# Apolipoprotein E4 Exacerbates Aβ Oligomer-Induced Synaptotoxicity through Promoting Autophagic Degradation of Glutathione Peroxidase 4

**DOI:** 10.64898/2026.11.01.763365

**Authors:** Jing Zhou, Hongji Liao, Wenjing Liang, Shuyi Liu, Jingjing Cai, Yu Liu, Yimei Liu, Zhonglou Zhou, Shengzhou Wu

**Affiliations:** State Key Laboratory of Eye Health, Eye Hospital, Wenzhou Medical University, Wenzhou, 325027, China

**Keywords:** Macroautophagy, Alzheimer, APOE4, STIM1, STIM2, SOCE, Bcl2, P62, Mitochondria

## Abstract

Extensive synaptic loss and progressive neurodegeneration are defining pathological hallmarks of Alzheimer’s disease (AD), primarily driven by soluble Aβ oligomers (AβO) and neurofibrillary tangles. The apolipoprotein E4 (APOE4) isoform further aggravates these pathological cascades, yet the precise molecular mechanisms underlying AD pathogenesis remain poorly elucidated. Store-operated calcium entry (SOCE) deficiency in neurons has been proven to exert detrimental effect on dendritic spine integrity and function. Herein, we investigated whether SOCE dysregulation contributes to AβO-induced synaptotoxicity and how APOE4 modulates this pathological process. In primary neuronal cultures, co-treatment with AβO and APOE4 markedly reduced synapse density and impaired dendritic arbor complexity, compared with AβO treatments combined with APOE3 or APOE2 isoforms. Consistently, neurons exposed to AβO plus APOE4 exhibited significantly lower SOCE amplitudes than those treated with AβO and APOE3. In line with SOCE suppression, the AβO/APOE4 combination specifically downregulated stromal interaction molecule 1 (STIM1) protein expression, with no observable change in STIM2 levels. Mechanistically, AβO/APOE4-mediated STIM1 reduction was dependent on p62-selective autophagic degradation. Notably, exogenous STIM1 overexpression effectively rescued the deficits in dendritic arbor complexity. Furthermore, STIM1 downregulation induced by AβO and APOE4 initiated the autophagic degradation of glutathione peroxidase 4 (Gpx4). This autophagy activation was attributed to decreased Bcl2 expression and occurs independently of STING accumulation. Subsequent Gpx4 insufficiency further triggered excessive accumulation of reactive oxygen species and disrupted mitochondrial cristae morphology in neurons. Collectively, our findings uncover a previous unrecognized mechanism whereby APOE4 exacerbates AβO-induced synaptotoxicity. Specifically, the combination facilitates p62-mediated selective autophagic degradation of STIM1, which suppresses Bcl2 expression and ultimately promotes robust Gpx4 autophagic degradation, driving neuronal synaptic damage.

**Graphic abstract:** 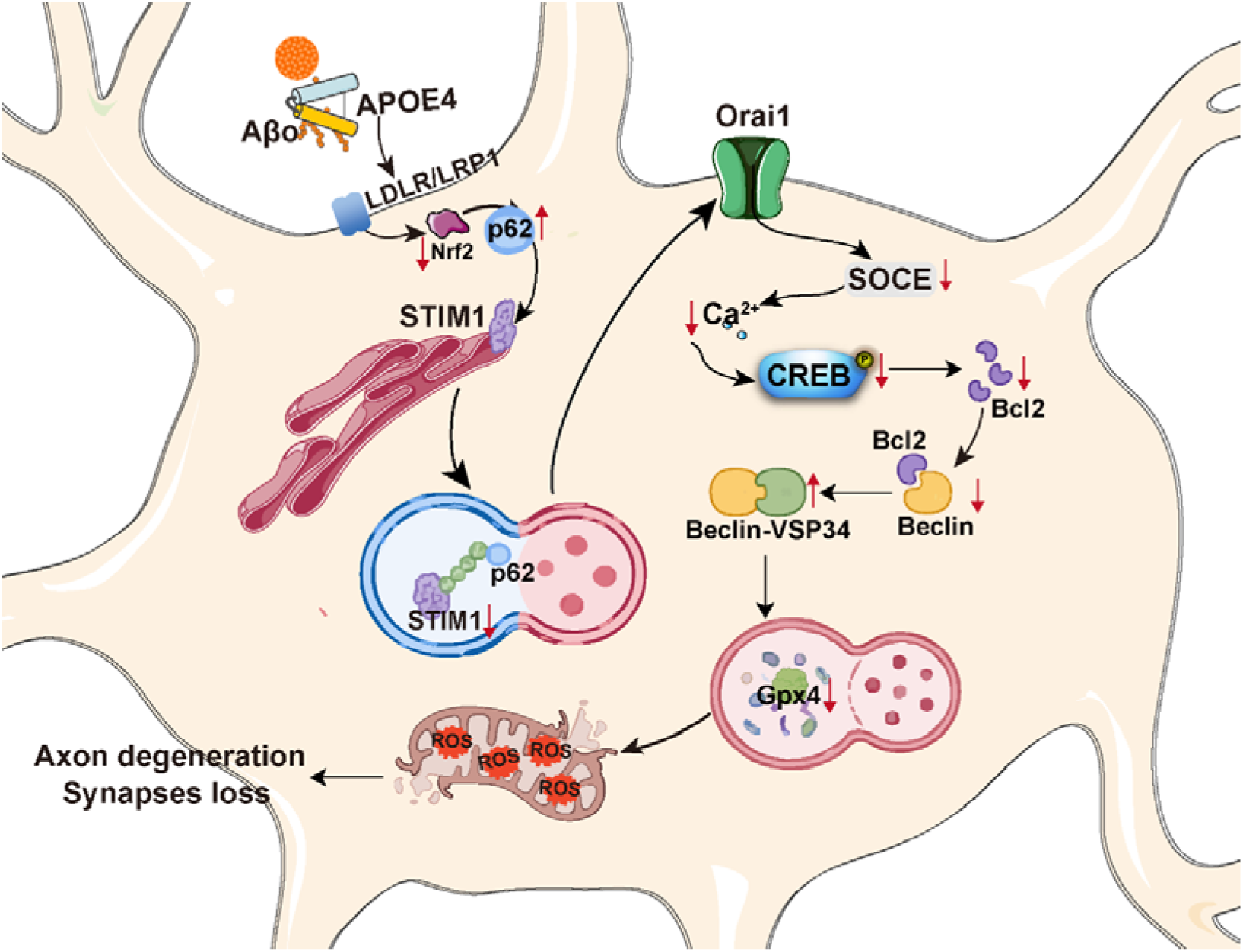

**Schematic overview of this study:** Aβ oligomers (Aβo), together with APOE4, suppress store-operated calcium entry (SOCE) via STIM1 protein downregulation, which reduces dendritic arbor complexity and compromises synaptic integrity. STIM1 depletion is mediated through the p62-dependent selective autophagy pathway. Deficient SOCE further lowers Bcl2 expression, thereby inducing bulk autophagy-dependent degradation of glutathione peroxidase 4 (GPX4) — a critical antioxidant enzyme. This cascade drives reactive oxygen species (ROS) accumulation and disrupts mitochondrial cristae architecture.

## Introduction

Synapse loss represents a core pathological hallmark of Alzheimer’s disease (AD). Temporally, amyloid-β (Aβ) deposition precedes synapse loss and tau aggregation during AD progression ^1^. In the AD brain, Aβ triggers synaptic dysfunction and neuroinflammation, and further promotes tau deposition in AD brain, whereas neurofibrillary tangles are strongly correlated with neurodegeneration and AD clinical manifestations ^1–3^. Aβ exerts prominent synaptic neurotoxicity. Specifically, soluble Aβ oligomers purified from the cerebral cortex of AD patients have been shown to reduce dendritic spine density and inhibits long-term potentiation in the rodent hippocampus ^4^. Multiple mechanisms have been proposed to underlie AD-associated synapse loss, including autophagy-mediated synaptic degradation and complement-dependent microglial synaptic pruning ^5, 6^. Nevertheless, the precise molecular mechanisms driving synapse loss and subsequent neurodegeneration in AD remains poorly defined.

Apolipoprotein E (APOE) is essential for the transport and clearance of Aβ. APOE polymorphism is widely acknowledged as the strongest genetic risk factor for late-onset AD. Carriers of the APOEε4 allele display accelerated and earlier AD onset, whereas the APOEε2 allele confers protective effects with reduced AD risk relative to the predominent APOEε3 allele ^7, 8^. Secreted APOE assembles into lipoprotein particles by associating with phospholipids and cholesterol transported by ATP-binding cassette transporters A1 and G1, thereby facilitating cerebral Aβ trafficking ^9^. By binding to surface receptors, including the low-density lipoprotein (LDL) receptor, LDL receptor-related protein 1, and triggering receptor expressed on myeloid cells 2 (TREM2) ^10, 11^, Aβ can be engulfed and internalized by astrocytes and microglia, followed by intracellular proteolytic degradation. Additionally, soluble Aβ is eliminated through the glymphatic system and the blood-brain barrier, both of which mediate Aβ clearance in an APOE isoform-dependent manner. Specifically, APOE4 exhibits markedly lower clearance efficiency than APOE2 and APOE3 ^12, 13^. Beyond regulating soluble Aβ clearance, APOE4 increases synaptic accumulation of soluble Aβ by over five-fold relative to APOE3 ^14^. In contrast, APOE3, but not APOE4, confers protection against soluble Aβ-triggered synaptic toxicity ^15^. Thus, APOE4 acts as a cofactor that exacerbates soluble Aβ-mediated synaptic toxicity by altering Aβ clearance and distribution. Nevertheless, the mechanism underlying direct modulation of Aβ synaptotoxicity by APOE4 remains poorly understood.

Store-operated calcium entry (SOCE) is a fundamental cellular mechanism responsible for maintaining intracellular basal calcium homeostasis. Depletion of luminal calcium store from the endoplasmic reticulum (ER) triggers the oligomerization of ER-resident stromal interaction molecules (STIM1 and STIM2), which subsequently relocate and tether to the plasma membrane. This conformational and positional rearrangement activates plasma membrane channels, including ORAI or transient receptor potential canonical (TRPC) channels, thereby driving extracellular calcium influx ^16, 17^. In the central nervous system, impaired SOCE has been widely documented in experimental AD models and postmortem human AD brain tissues. For instance, compromised neuronal SOCE destabilizes neuronal synaptic structure in presenilin1 knock-in mice, downregulated STIM2 expression has been detected in the cortical tissues of AD patients ^18, 19^. Notably, STIM1 levels in postmortem AD brains at Braak stage VI are approximately 70% lower than those in non-demented control subjects ^20^. Additionally, STIM1B—a proteolytic cleavage variants of STIM1 localized to presynaptic ER compartments—is significantly reduced in AD patients, and this deficiency impairs Orai1 channel-dependent short-term synaptic plasticity ^21^. Given that compromised neuronal SOCE is associated with unstable neuronal synapse and simplified dendrite arbors ^5, 18^, the present study investigates the regulatory role of SOCE in AβO-induced synaptotoxicity, as well as the modulatory effect of APOE4 on this pathological process.

## Materials and Methods

### Reagents and Tools table

| **Reagents/ resources** | **Reference or source** | **Identifier or catalog numbers** |
|---|---|---|
| **Experimental models** |  |  |
| HEK-293 cells (*H. Sapiens*) | ATCC | CRL-1573 |
| C57BL6/J (*M. musculus*) | Shanghai Animal Experimental Center, Chinese Academy of Sciences | N/A |
| **Recombinant DNA** |  |  |
| pEX-SP-EYFP-STIM1(D76A) | Liou *et al* (2005) | Addgene (cat#18859) |
| pEX-SP-EYFP | This study | N/A |
| ptfLC | Kimura *et al*(2007) | Addgene(cat#21074) |
| pUC19-hNFE2L2 | Sino Biological | cat#HG17384-U |
| pUC19 | Sino Biological | cat#D2602 |
| **Antibodies** |  |  |
| STIM1 | Cell Signaling Technology | RRID#AB_2271287/cat#4916 |
| STIM2 | Cell Signaling Technology | cat#4917, RRID#AB_2198021 |
| SQSTM1/p62 | Cell Signaling Technology | cat#5114, RRID#AB_10624872 |
| α-tubulin | Cell Signaling Technology | cat#2144, RRID#AB_2210548 |
| phospho–cofilin (Ser3) | Cell Signaling Technology | cat#3313, RRID#AB_2080597 |
| phospho-LIMK1(Thr508)/LIMK2 (Thr505) | Cell Signaling Technology | cat#3841 |
| LC3B | Cell Signaling Technology | cat#2775, RRID#AB_915950 |
| TFEB | Cell Signaling Technology | cat#37785, RRID#AB_2799119 |
| Glutathion peroxidase 4 (Gpx4) | Proteintech | cat#67763-1-Ig, RRID#AB_2909469 |
| MAP2 | Proteintech | cat#67015-1-lg, RRID#AB_2882331 |
| Nrf2 | Proteintech | cat#16396-1-AP, RRID#AB_2782956 |
| Synapsin I | Millipore | cat#AB1543P |
|  |  | RRID#AB_90757 |
| STING | ABclonal | cat#A21051<br>RRID# AB_3083450 |
| Bcl2 | Abcam | cat#ab182858<br>RRID# AB_2715467 |
| Horseradish peroxidase (HRP)-conjugated goat anti-rabbit IgG (H + L) | Zhongshan Golden Bridge Biotechnology | cat#ZB-2301 |
| anti-mouse IgG (H + L) | Zhongshan Golden Bridge Biotechnology | cat# ZB-2305 |
| **Oligonucleotides and sequence-based reagent** |  |  |
| PCR primers | This study(Sangon Biotech) | N/A |
| **Chemicals, enzymes and other reagents** |  |  |
| Neurobasal<sup>®</sup> medium | Gibco | cat#21103–049 |
| DMEM | Gibco | cat#11995 |
| B27 | Gibco | cat#17504-044 |
| GlutaMAX | Gibco | cat#35050-061 |
| Fetal bovine serum | BioInd | cat#04-001-1A |
| Human APOE2 | PeproTech | cat#350-12 |
| Human APOE3 | PeproTech | cat#350-02 |
| Human APOE4 | PeproTech | cat#350-2-04 |
| TRIzol reagent | Invitrogen | cat #15596-018 |
| HiScript II 1st strand cDNA synthesis kit | Vazyme | cat#R211 |
| Taq Pro Universal SYBR qPCR master mix | Vazyme | cat#Q712 |
| NaF | SINOPHARM | cat#10019608 |
| Na3VO4 | Sigma | cat#S6508-10G |
| Protease inhibitor cocktails | Sigma | cat#p8340 |
| Polyethyleneimine | Sigma | cat#P3143 |
| Dimethyl sulfoxide | Sigma | cat#D8418 |
| Phenylmethanesulfonyl fluoride (PMSF) | Beyotime Biotechnology | cat#ST506 |
| Penicillin–streptomycin | Beyotime Biotechnology | cat#C0222 |
| DL-dithiothreitol | Beyotime Biotechnology | cat#ST041 |
| the enhanced BCA protein assay kit | Beyotime Biotechnology | cat#P0010 |
| human Aβ peptide (1-42) | Beyotime Biotechnology | cat#P9001 |
| Nitrocellulose (NC) membranes | Cytiva | 0.22 µm, cat#10600001; 0.45 µm, cat#10600002 |
| SuperSignal<sup>®</sup> West Femto maximum sensitivity substrates | ThermoFisher | cat#34095 |
| Lipofectamine 3000 | Invitrogen | cat#L3000015 |
| Fura-2 AM | Invitrogen | cat#F1221 |
| pluronic F127 | Invitrogen | cat#P6867 |
| 2'-7' Dichlorofluorescin diacetate (DCFH-DA) | Dojindo | cat#R252 |
| **Software** |  |  |
| GraphPad Prism 8.0 | https://www.graphpad.com |  |
| SPSS 23.0 | https://www.ibm.com/cn-zh/products/spss-statistics |  |
| ImageJ | https://imagej.net/software/fiji/downloads |  |

### Plasmid construction and transfection

The original plasmid pEX-SP-EYFP-STIM1(D76A) was developed by Tobias Meyer’s group ^22^ and served as a template for constructing the control plasmid, pEX-SP-EYFP. The EYFP fragment (740 bp) was amplified using a PCR reaction with the forward primer GACCGGTCGCCACCATGGTGAG, which incorporated the AgeI restriction site, a Kozak consensus sequence, and an ATG initiator codon. The reverse primer, GAAGATCTCTACTTGTACAGCTCGTCCATGCCG, included the BglII restriction site and a TAG terminator codon. After digesting both pEX-SP-EYFP-STIM1(D76A) and the PCR product, ligation was performed, resulting in the replacement of EYFP-STIM1 with the PCR product, EYFP, in the control plasmid, pEX-SP-EYFP. The EYFP sequences were subsequently verified through sequencing. HEK293 cells were cultured at a density of 50,000 cells per 24-well dish, while primary hippocampal neuronal cultures were maintained at a density of 25,000 cells per 96-well dish. After 24 h, the HEK293 cells were transfected with 500 ng of the pUC19-hNFE2L2 plasmid using 0.5 μL of Lipofectamine 3000 per 24-well dish to overexpress Nrf2, or with an equal amount of the control pUC19 plasmid, for a duration of 2 to 3 days. At DIV 4-5, the primary neurons were transfected with 100 ng of the pEX-SP-EYFP-STIM1(D76A) plasmid using 0.25 μL of Lipofectamine 3000 per 96-well dish to overexpress STIM1(D76A) or the control plasmid, pEX-SP-EYFP, for 2 days.

### Neuronal cultures

Hippocampal neuronal cultures were established from neonatal C57BL/6J mice on day 1. The pups were euthanized on ice and subsequently decapitated. The skulls and meninges were removed, and the hippocampal tissues were isolated, cut into small pieces, and digested in 0.25% trypsin/EDTA at 37℃ for 10 min. The tissues were gently triturated through a fire-polished glass pipette. The resulting suspension was plated in polyethyleneimine-coated 96-well or 24-well culture dishes containing DMEM supplemented with 10% FBS. After 4 to 6 h, the media were replaced with neurobasal medium containing 2% B27, and 0.2 mM L-glutamine, and a 1% penicillin-streptomycin (PS) mixture. After 48 h, a mitotic inhibitor, cytosine arabinoside (1-2 μM), was added to the cultures for 5 to 7 days. Following the removal of the inhibitor, the cultures were utilized on days *in vitro* (DIV) 7 to 9 days. HEK293 cells were cultured in DMEM supplemented with 10% FBS and a 1% PS mixture. All cells were maintained in a 37°C incubator with 5% CO2.

### Human AβO preparation

Human Aβ1-42 oligomers were prepared as previously described [28]. Briefly, Aβ1-42 powder was dissolved in 1,1,1,3,3,3-hexafluoro-2-propanol and stored at 4℃ for 30 min. The solution was then dried at room temperature, and the resulting dry pellet was reconstituted in DMSO to achieve a concentration of 1 mM. The peptide was subsequently diluted to 500 nM in phenol-free F12 medium and incubated at 4℃ for 24 h. The peptide solution was centrifuged at 14,000 × g for 30 min at 4°C, and the supernatant was collected as Aβ1-42 oligomers.

### SiRNA knockdown experiment

Primary hippocampal neurons at DIV 5–7 were transfected with siRNA using Lipofectamine 3000 in a 96-well plate. According to the manufacturer’s instructions: Solution A was prepared by mixing 0.3 μL Lipofectamine 3000 with 5 μL Opti-MEM per well. Solution B was prepared by mixing 20 pmol siRNA with 5 μL Opti-MEM per well. sip62 sequences: sense 5‘-GGUUGACAUUGAUGUGGAATT-3’, antisense 5‘-UUCCACAUCAAUGUCAACCTT-3’. siSTING sequences: sense 5‘-GAGCAG GCUAAACUCUUCUTT-3’, antisense 5‘-AGAAGAGUUUAGCCUGCUCTT-3’. siNC sequences: sense 5‘-UUCUCCGAACGUGUCACGUTT-3’, antisense 5‘-ACGUGACACGUUCGGAGAATT-3’. Solution A and Solution B were combined to form Solution C, followed by incubation at room temperature for 10–15 min. For cell treatment, 100 μL neuronal culture medium was added to each well, and then 15 μL of Solution C was supplemented per well. Subsequent experiments were performed 48 hours after transfection of siRNA.

### Western blotting

Primary neuronal cultures or HEK293 cell cultures were washed twice with PBS and lysed on ice using Super RIPA buffer, which contained 500 mM Tris base, 150 mM NaCl, 1 mM Na2EDTA, 1% NP-40, a 1× protease inhibitor cocktail, 1 mM phenylmethanesulfonyl fluoride, 1 mM dithiothreitol, 1 mM sodium vanadate, and 1 mM sodium fluoride. Equal amounts of protein (20-40 µg) were resolved by 12% SDS-PAGE and transferred to a nitrocellulose membrane. Following this procedure, the membranes were blocked with 5% skim milk and subsequently incubated with a primary antibody (1:500 or 1:1000) at 4℃ overnight. After washing with PBS-T (PBS containing 0.1% Tween-20), the membranes were incubated with a peroxidase-conjugated anti-mouse or anti-rabbit secondary antibody (1:2000) for 1 h at room temperature. Blots were detected using enhanced chemiluminescence detection reagents. The density of the bands was quantified using ImageJ software (National Institutes of Health, USA) and normalized to α-tubulin.

### Immunofluorescence

The primary hippocampal neurons were plated on coverslips in 24-well plates. After treatment, the neurons were washed twice with PBS and fixed with 4% paraformaldehyde at room temperature for 20 min. The cells were then permeabilized with 0.1% Triton X-100 in PBS at room temperature for 30 min. Neurons were blocked with 10% goat serum in PBS for 20 min at room temperature and subsequently incubated with a primary antibody (1:300) in PBS containing 0.1% Triton X-100 at 4℃ overnight. After washing with PBS twice, the coverslips were incubated with donkey anti-rabbit IgG (H+L) Alexa Fluor® 594 secondary antibody for 1 h at room temperature. Following washing, the slides were mounted with mounting media containing DAPI to label the cell nuclei. Images were acquired using a confocal fluorescence microscope system (Carl Zeiss, Germany) with a 20× objective.

### Examination of reactive oxygen species

The detection of reactive oxygen species (ROS) in primary neurons was conducted on DIV 7-9. Cells were treated with APOE and AβO for 24 h, and then incubated with the ROS fluorescent probe DCFH-DA at 37°C in the dark for 30 min. Images were captured using a 20× objective on a DMi8 fluorescence microscope (Leica, Germany). The fluorescence intensity of DCFH-DA was analyzed using ImageJ software (National Institutes of Health, USA).

### RNA extraction and real-time quantitative PCR

Total RNA was extracted from primary neurons or HEK293 cells using Trizol. The RNA concentration was measured with a NanoDrop 2000 spectrophotometer (Thermo Fisher, USA). cDNA was synthesized using 1000 ng of RNA as a template with the HiScript II 1st Strand cDNA Synthesis Kit, following the manufacturer’s instructions. The PCR reaction was performed using Taq Pro Universal SYBR qPCR Master Mix in the QuantStudio 5 system (ABI, USA), with GAPDH serving as the internal reference. The primers used in qPCR were as follows: mouse GAPDH, forward: GAAGGTCGGTGTGAACGGATTTGG, reverse: TGTAGACCATGT AGTTGAGGTCA; human GAPDH, forward: ATGGGGAAGGTGAAGGTCG, reverse: GGGGTCATTGATGGCAACAATA; mouse STIM1, forward: CAGGG ACTGTACTGAAGATGACA, reverse: AGGTGATTATGCCGAGTCAAG; human STIM1, forward: TGTGGAGCTGCCTCAGTATG, reverse: CTTCAGCACAGTCCCTGTCA; mouse STIM2, forward: TCGAAGTGGACGAGAGTGATG, reverse: TTTCCACTGTTTCCACAAATCC; human Nrf2, forward: TTCCCGGTCACATCGAGAG, reverse: TCCTGTTGCATACCGTCTAAATC. The relative levels of target genes were analyzed using the comparative 2^−ΔΔCT^ method.

### Measurement of SOCE and intracellular calcium with a ratiometric calcium imaging

Following our established protocol, calcium influx through store-operated calcium channels was measured using Fura-2, a fluorescent dye for intracellular calcium detection [33, 34]. Briefly, primary neurons were incubated in a loading buffer containing 144 mM NaCl, 10 mM glucose, 10 mM HEPES, 5 mM KCl, 1.8 mM CaCl2, 1 mM MgCl2, 0.01 mM glycine, 2 μM Fura-2 AM, and 0.05% pluronic acid, adjusted to pH 7.2, in the dark at 37°C for 45 min. The neurons were then washed in a dye-free loading solution for an additional 45 min in the dark at 37℃. Prior to determining the calcium influx, the solution was replaced with a calcium-free solution containing EGTA. Thapsigargin (5 μM) was added to inhibit ER Ca^2+^-ATPase (SERCA) and induced calcium depletion from the endoplasmic reticulum. Calcium (1.8mM) was then added to the medium, resulting in a significant increase of intracellular calcium concentration [Ca^2+^]_i_, indicative of SOCE. The detection system comprised an EMCCD camera (Andor, UK) and a high-speed monochromator (Photon Technology International, USA). Images were acquired using a 40x objective on an inverted microscope (Nikon, Japan). Thapsigargin also triggers store-independent calcium entry in endothelial cells via activation of the PLC/PKC pathway ^23^, an effect not investigated in the present study.

### Autophagy flux assay

Primary neuronal cultures were plated on coverslips sitting in 24-well culture plates. At DIV 4-5, the cells were transfected for 48 h in Opti-MEM medium using Lipofectamine 3000 ^TM^ with 500 ng of mRFP-GFP tandem LC3 (ptfLC3) plasmid (Addgene) ^24^. After transfection, the neruons were treated with AβO (200 nM) in conjunction with different isoforms of APOE (200 nM for each) for 24 h. The neurons were fixed with 4% formaldehyde in PBS for 12 min. After being washed with PBS three times, cells were mounted on glass slides with mounting medium containing 4,6-diamidino-2-phenylindole (DAPI) and observed under a Zeiss LSM 880 confocal microscope system (Carl Zeiss), under 63×objective. The images were processed with Zen Le software (Carl Zeiss).

### Transmission electron microscopy

Neurons at a density of 1×10^4^ neurons were seeded on 24-well plates, and treated with APOE2+AβO or APOE4+AβO for 24 h. Neurons were harvested by trypsinization and centrifuged. The cell pellets were fixed with 2.5% glutaraldehyde fixative (pH 7.0-7.5) at 4℃ for 2–4 h. Subsequent embedding, sectioning, mounting, and transmission electron microscope (TEM) imaging were performed by specialists in Servicebio electron microscope core facility (Servicebio, Wuhan, China). In brief, after discarding supernatant, cell pellets were pre-embedded in 1% agarose. Agarose-embedded samples were further fixed with 1% osmium tetraoxide (OsO_4_) for 2 h at room temperature, then dehydrated in a graded series of ethanol (30-100%) followed by acetone at room temperature. Dehydrated samples were embedded in EMBed812 and incubated overnight at 37℃, followed by polymerization in 60℃ for 48 h. Blocks were cut into 60–80nm ultrathin sections using a Leica UC7 ultramicrotome (Leica, Germany). Ultrathin sections were stained with 2% uranyl acetate and 2.6% lead citrate. Sample were observed and imaged under an HT7800 transmission electron microscope (Hitachi, Japan) at an accelerating voltage of 80 kV and 2500 ×magnification.

### Experimental Design and Statistical Analysis

Sample sizes were determined based on previous literature, and formal power calculations were not performed, as the study aimed to explore preliminary trends in different treatments. SPSS version 23.0 (IBM) software was utilized to conduct statistical analyses. To enable inter-experimental normalization across independent blots, densitometric ratios (target protein/α-tubulin) from all experiments were pooled per treatment group, and comparisons were made using these pooled values. Results were presented as means ± SEM. Data were assessed for normality using the Shapiro-Wilk test. If the data exhibited a normal distribution, parametric analyses were employed; otherwise, nonparametric analyses were conducted to compare differences. For parametric data, Student’s t-test was used to compare differences between two groups. Data from more than two groups were analyzed using one-way ANOVA, followed by the Bonferroni post hoc test. For nonparametric data, the Mann–Whitney U test was employed to compare differences between two groups, while the Kruskal–Wallis test was used to compare differences among multiple groups. A difference was considered statistically significant when p < 0.05.

## Results

### Defective SOCE mediated neuronal dendritic impairment induced by combined AβO and APOE4 treatment

To investigate the synergistic neurotoxic effects of AβO and APOE4 on neurites, we analyzed presynaptic protein levels following their individual and combined treatments. Treatment with 25, 50, or 100 nM AβO alone for 24 h did not reduce synapsin I protein levels in primary hippocampal neuronal cultures relative to vehicle control. However, when co-administration with 200 nM APOE4, 50 nM AβO significantly reduced synapsin I compared with AβO ( 50 nM) treatment alone, whereas the other two AβO doses not exerted such effect (Figure 1A). Given that 25 and 100 nM AβO for 24-h treatment exerted no effect on synapsin I protein levels, we raised AβO concentration to 200 nM for subsequent experiments. This adjustment was based on our prior work showing that 100 nM AβO elicits robust toxicity to axons and dendrites in primary neuronal cultures ^25^. The morphology and synaptic structure influenced by AβO and the different APOE isoforms were further examined. The primary hippocampal neuronal cultures were treated with AβO (200 nM) supplemented with equal amounts of APOE2, APOE3, and APOE4 (200 nM each) for 24 h. Following treatment, the cultures were stained for microtubule-associated protein 2 (MAP2) to evaluate neuronal dendritic arborization. Compared to AβO supplemented with either APOE2 or APOE3, dendritic arborization under combined AβO and APOE4, as revealed by MAP2 staining, was significantly impaired (Figure 1B). To assess the synapses, we labeled them in the cultures using the presynaptic marker synapse I, in conjunction with MAP2 staining. The synapse density was significantly reduced in the neuronal cultures treated with combined AβO and APOE4, compared to those treated with combined AβO and APOE3 (Figure 1C).

**Figure 1.**
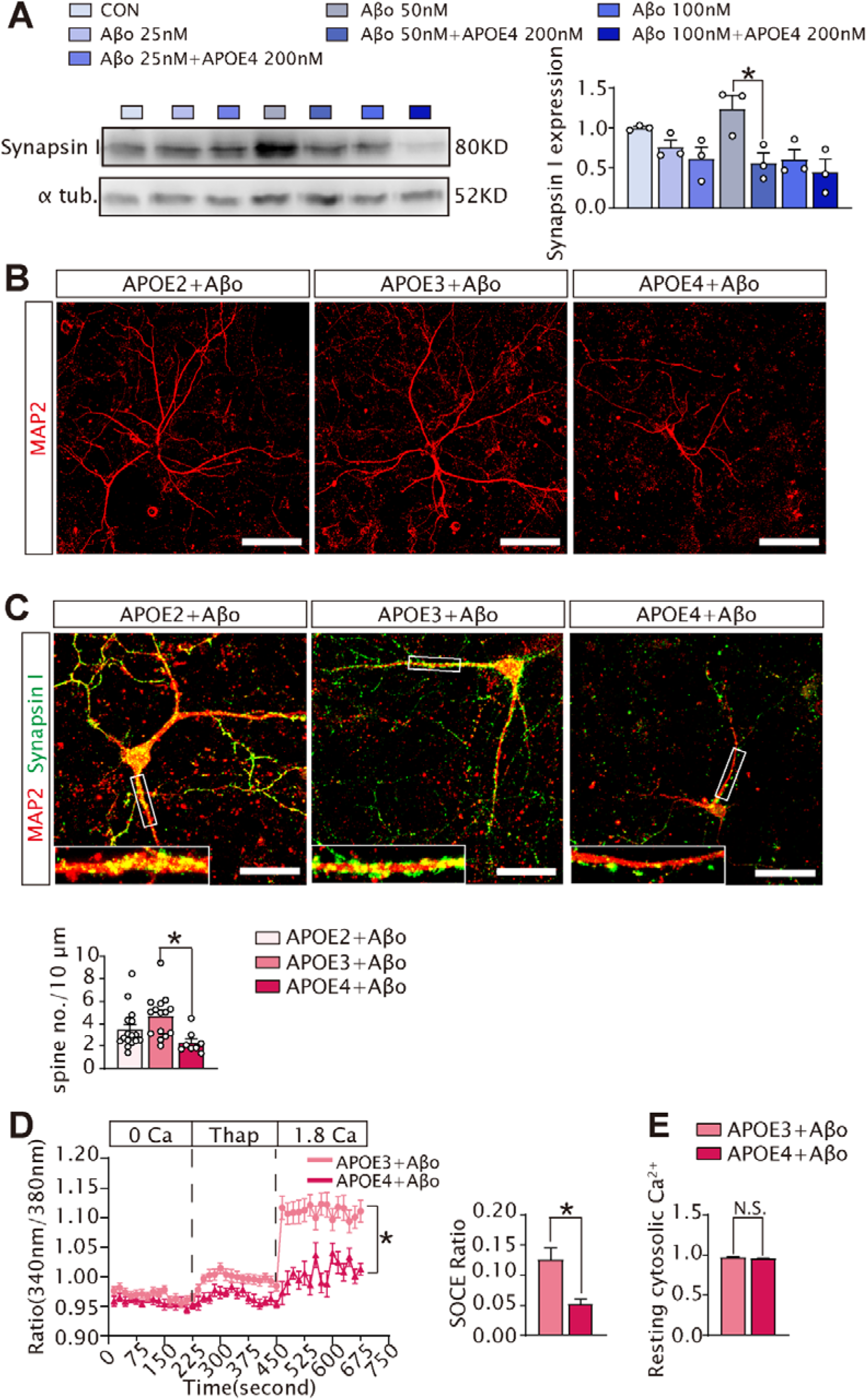
APOE4 enhanced synaptotoxicity in primary hippocampal neuronal cultures, an effect associated with downregulated SOCE activity. (A) Primary hippocampal neuronal cultures at days *in vitro* (DIV) 7-9 were treated with AβO (25, 50,100 nM), or along with APOE4 (200 nM each) for 24 h. Western blot was conducted to examine synapsin I protein levels in the protein lysates. Synapsin I protein levels in neurons treated with APOE4 + AβO (50 nM) was significantly lower than those in cells treated with AβO (50 nM) alone. One-way ANOVA, followed by Bonferroni post-hoc test, was employed for comparisons, F(6, 14) = 4.720, p = 0.008. Specifically, APOE4 + AβO (25 nM) *vs*. AβO(25 nM), p = 1.000; APOE4 + AβO(50 nM) *vs*. AβO(50 nM), p = 0.048; APOE4 + AβO (100 nM) *vs*. AβO(100 nM), p = 1.000. Primary hippocampal neuronal cultures were treated with AβO (200nM), along with APOE2, APOE3, or APOE4 (200 nM each) for 24 h. (B) These cultures were fixed and stained for a neuronal marker MAP2 (red). These images are typical of 12-14 neurons for each treatment, obtained from two separate slides. Images were acquired using a Zeiss LSM880 confocal laser microscope system with a 20× objective. Bar, 100μm for all panels. (C) The fixed cultures were co-stained for MAP2 (red) and a synaptic marker synapsin I (green). These images are representative of three slides containing 8-15 neurons for each treatment. Images were acquired with a 40×objective. Bar, 50 μm for all panels. Spine no./10 μm in axon in neurons treated with APOE4+ AβO were lower than that in APOE3+ AβO-treated neurons. Kruskal-Wallis test, was employed for comparisons, p = 0.002. Specifically, the results revealed significant differences: APOE2 + AβO vs. APOE4 + AβO, p = 0.199; APOE3 + AβO vs. APOE4 + AβO, p = 0.001. (D) After treatment, neurons were loaded with a ratiometric dye Fura-2 AM and subjected to calcium measurement in calcium-free buffer at initial measurement (0 Ca). After baseline calcium levels were obtained, calcium levels were measured under addition of an Ca^2+^-ATPase inhibitor, thapsigargin (5 μM) (Thap). At the final measurement, a regular buffer containing 1.8 mM Ca^2+^ was added into the cultures (1.8 Ca). Trend lines for calcium ratios are averaged from 21 to 29 neurons per time point of 3 independent cell cultures. (E) Mann-Whitney U test was employed to compare the differences in the SOCE amplitude, yielding a p value of 0.011. Additionally, a Student’s t-test was conducted to assess the differences in resting cytosolic free Ca2+ concentration, resulting in t(44.690)=1.596, p = 0.118.

SOCE is essential for dendritic stability ^18, 26^. Since combined AβO and APOE4 negatively affected dendritic arborization and decreased synapse density (Figures 1B and 1C), we investigated whether AβO, when paired with APOE4, modulated SOCE in neurons. After a 24-h treatment with AβO (200 nM) and APOE4 or APOE3 (200 nM), neurons were loaded with the ratiometric dye Fura-2 AM. The emission fluorescence ratios under 340/380 nm excitation wavelengths were recorded to assess calcium levels in the neurons. Under normal calcium conditions, intracellular calcium levels ([Ca^2+^]_i_) were measured as resting cytosolic free calcium. Compared to the neurons treated with combined AβO and APOE3, the SOCE amplitudes in the neurons treated with combined AβO and APOE4 were significantly reduced (Figure 1D). However, the baseline [Ca^2+^]_i_ levels in neurons treated with AβO and APOE4 were similar to those in neurons treated with AβO and APOE3 (Figure 1E).

### Combined AβO and APOE4 treatment downregulated STIM1 protein

Accumulated studies suggest that the downregulation of STIM1 or STIM2 may contribute to synaptic deficits observed in AD. Thus, we began to examine the effects on STIM1 and STIM2 under the combined treatment of AβO with different APOE isoforms. We reprobed the membranes for Figure 1A and observed that treatment with 25, 50, or 100 nM AβO alone exerted no obvious effect on the protein levels of STIM1 or STIM2. By contrast, combined treatment with APOE4 (200 nM) and AβO (50 nM) markedly downregulated STIM1 protein levels relative to AβO (50 nM) alone, and this synergistic inhibitory effect was not detected for STIM2 (Figure 2A). To probe dose-response effect, co-administration of 200 nM AβO and 200 nM APOE4 produced a significantly more pronounced reduction in STIM1 expression compared with 100 nM AβO plus 100 nM APOE4 or 400 nM AβO plus 400 nM APOE4 (Supplemental Figure 1). We further compared the synergistic effects of different APOE isoforms combined with 200 nM AβO on STIM1 and STIM2 by measuring the mRNA and protein levels of both genes in neuronal cultures. Combined treatment with 200 nM AβO and 200 nM APOE4 markedly reduced STIM1 protein levels compared with co-administration of 200 nM AβO plus either 200 nM APOE2 or APOE3. Notably, STIM1 mRNA levels remained comparable across the three treatment groups (Figures 2B and 2C). By contrast, neither STIM2 mRNA nor protein levels exhibited significant differences among all the experimental groups (Figures 2C and 2D).

**Figure 2.**
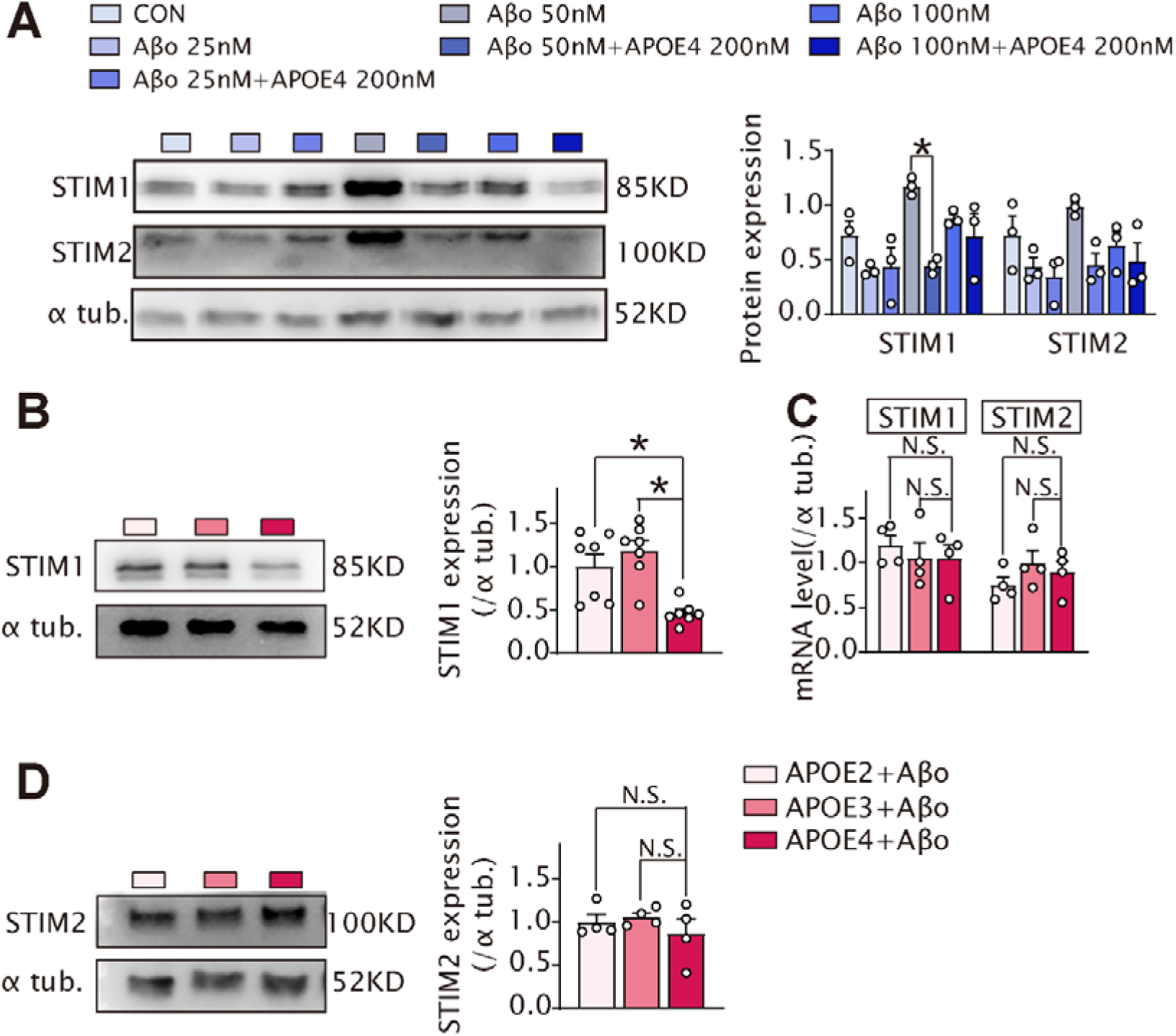
APOE4 combined with AβO reduced STIM1 protein levels, but had no significant effect on STIM2. **(A)** Primary hippocampal neuronal cultures at DIV7-9 were subjected to treatments with 25, 50, or 100 nM AβO, in combination with 200 nM APOE4. Western bot was used to examine STIM1 and STIM2 protein levels. STIM1 protein levels in neurons treated with APOE4 + AβO (50 nM) were significantly lower than those in cells with treated with AβO (50 nM) alone. One-way ANOVA, followed by Tamhane T2 test, was employed for comparisons, F(6, 14) = 5.817, p = 0.003. Specifically, APOE4 + AβO (25 nM) *vs*. AβO(25 nM), p = 1.000; APOE4 + AβO (50 nM) *vs*. AβO (50 nM), p = 0.007; APOE4 + AβO (100 nM) *vs*. AβO (100 nM), p = 1.000. There was no significant difference in STIM2 protein between APOE4 + AβO treatment and AβO treatment alone. One-way ANOVA, followed by Bonferroni post-hoc test, was employed for comparisons, F(6, 14) = 2.933, p = 0.046. Specifically, APOE4 + AβO (25 nM) *vs*. AβO(25 nM), p = 1.000; APOE4 + AβO (50 nM) *vs*. AβO (50 nM), p = 0.229; APOE4 + AβO (100 nM) *vs*. AβO (100 nM), p = 1.000. Primary hippocampal neuronal cultures were subjected to treatments with AβO (200 nM), in conjunction with either APOE2, APOE3, or APOE4 (200 nM each) for 24 h. Seven independent cell cultures were prepared for each treatment. (**B**) Western blot was conducted to analyze STIM1 protein levels in the protein lysates. One-way ANOVA, followed by Tukey test, was employed for comparisons. The results indicated F(2, 18) = 10.715, p = 0.001. Specifically, the results revealed significant differences: APOE2 + AβO *vs*. APOE4 + AβO, p = 0.030; APOE3 + AβO *vs*. APOE4 + AβO, p = 0.002. (**C**) RNA was extracted from the treated neurons and RT-qPCR was performed. Four independent cell cultures were prepared for each treatment. One-way ANOVA, followed by Bonferroni test, was used to compare the differences in STIM1 and STIM2 mRNA. The results indicated F(2,9) = 0.336, p =0.723 for STIM1 mRNA comparisons, and F(2,9) = 1.092, p = 0.376 for STIM2 mRNA comparisons. (**D**) Western blot was conducted to assess STIM2 protein levels. Kruskal-Wallis test was employed to compare the differences in STIM2 levels among various treatments, yielding p = 0.584.

### Downregulation of STIM1 by combined AβO and APOE4 treatment depended on p62-selective autophagic degradation

In numerous studies, AβO has been shown to induce oxidative stress in neurons. Compared to combined AβO and APOE2 or APOE3, levels of reactive oxygen species (ROS) were significantly higher in neurons treated with AβO and APOE4 (Figure 3A), while the levels of nuclear factor erythroid 2-related factor 2 (Nrf2) were significantly lower (Figure 3B). Nrf2 modulates protein abundance through both transcriptional and non-transcriptional mechanisms. At the transcriptional level, Nrf2 acts as a master transcription factor that binds to antioxidant response elements (AREs) within the promoter regions of target genes, thereby conferring protection against oxidative stress ^27^. At the non-transcriptional level, SQSTM1/p62 competes with Nrf2 for KEAP1 binding, forming the canonical p62-KEAP1-Nrf2 pathway, that promotes selective autophagy to mitigate oxidative stress ^28, 29^. Furthermore, oxidative stresses upregulate p62 to potentiate autophagy, sustaining cellular survival under oxidative challenge ^30^. Accordingly, we propose that the downregulation of STIM1 mediated by AβO and APOE4 is likely dependent on p62-mediated selective autophagy. Indeed, knockdown of p62 reversed STIM1 degradation induced by combined AβO and APOE4 treatment (Figure 3C). By contrast, knockdown of p62 did not alter STIM2 levels upon combined AβO and APOE4 treatment (Figure 3C).

**Figure 3.**
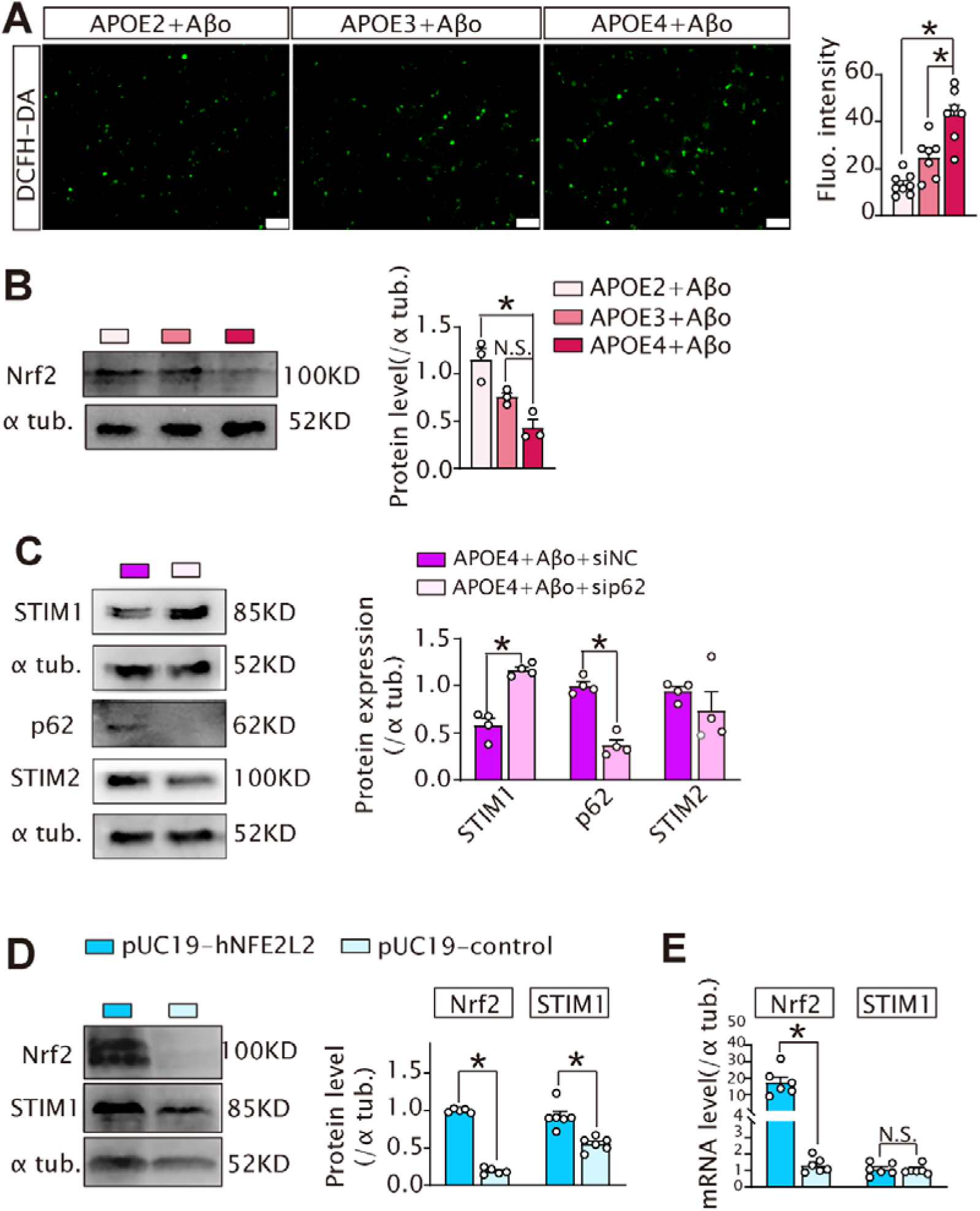
APOE4 together with with AβO downregulated STIM1; this effect was depended on p62-selective autophagic degradation. Primary hippocampal neuronal cultures at DIV7-9 were subjected to treatments as indicated. (**A**) ROS was indicated with a DCFH-DA probe. The levels were analyzed with One-way ANOVA, followed by Bonferroni test, F (2, 19) = 23.458, p < 0.0001. The ROS levels in neurons treated by APOE4 + AβO (n = 7) were either higher than those in APOE2 + AβO (n = 8) (p < 0.0001), or higher than those in APOE3 + AβO (n = 7) (p = 0.002). Images were acquired with DMi8 fluorescence microscope (Leica) under 20×objective. Bar, 100 μm for all panels. (**B**) Nrf2 protein levels were also examined in the treated neuron. One-way ANOVA, followed by Bonferroni test, was used for comparison, F (2, 6) = 16.058, p =0.004. Specifically, Nrf2 levels treated with APOE4 + AβO (n = 3) were significantly lower than those treated with APOE2 + AβO (n = 3)(p = 0.004), whereas Nrf2 levels treated with APOE4 + AβO were not different with those treated with APOE3 + AβO (n = 3) (p = 0.129). (**C**) Primary neuronal cultures were transfected with p62-specific siRNA, or scrambled siRNA (siNC) with lipofectamine 3000^TM^. After 48 h, the transfected neurons were treated with APOE4+ AβO for 24 h. STIM1, p62, and STIM2 protein levels were examined by western blot. Student’s *t-test* was used for comparing the differences of all protein levels. STIM1 levels in neurons transfected with the sip62 (n = 4) were higher than those transfected with siNC (n = 4), t(6) = 7.371, p < 0.0001. P62 levels in neurons transfected with the sip62 (n = 4) were lower than those transfected with siNC (n = 4), t(6) = 8.714, p < 0.0001. STIM2 in neurons transfected with the sip62 (n = 4) were not significantly different with those transfected with siNC (n = 4), t(6) = 1.015, p =0.349. (**D**) HEK293 cells were transfected with a pUC19-hNFE2L2 plasmid to overexpress Nrf2, or pUC19 control plasmid with lipofectamine 3000^TM^ for 48 h. Nrf2 and STIM1 protein levels were examined by western blot. (**E**) Student’s *t-test* was used for comparing the differences of either Nrf2 or STIM1 levels. Nrf2 levels in HEK293 cells transfected with the pUC19-hNFE2L2 plasmid (n = 5) were higher than those transfected with control plasmid (n = 5), t(8) = 37.374, p < 0.0001. STIM1 levels were also increased, t(10) = 4.740, p =0.001. Nrf2 and STIM1 mRNA were examined with RT-qPCR. Compared to those in cells transfected with control plasmid (n = 6), Nrf2 mRNA levels in cells overexpressing Nrf2 were higher, t(10) = 7.750, p = 0.001, whereas STIM1 mRNA levels were not different, p=1.000. Student’s *t-test* was used for comparing differences of Nrf2 mRNA, and Mann-Whitney U test for comparing differences of STIM1 mRNA.

To further substantiate that the p62-KEAP1-Nrf2 pathway modulates STIM1 abundance, we examined STIM1 levels in HEK293 cells with ectopic Nrf2 overexpression. In comparison to cells transfected with a control plasmid, both protein and mRNA levels of Nrf2 were significantly elevated in cells overexpressing Nrf2 (Figures 3D and 3E). STIM1 protein levels were significantly increased in HEK293 cells overexpressing Nrf2; however, the mRNA levels of STIM1 did not differ from those in cells transfected with the control plasmid (Figures 3D and 3E). Thus, p62-dependent autophagic degradation modulates STIM1 levels.

### Overexpression of STIM1 ameliorated neurite deficits induced by APOE4 and AβO

Cofilin is a key regulator of filamentous actin (F-actin) assembly and disassembly dynamics; dephosphorylation-mediated cofilin activation destabilizes dendrites and dendritic spines ^31^. Our prior work indicates that endoplasmic reticulum stress triggers autophagy, which promotes STIM2 degradation cofilin dephosphorylation, ultimately causing dendritic arbor disruption (33). To explore whether STIM1 depletion drives synapse loss and dendritic simplification via a comparable mechanism, we assessed phosphorylation status of cofilin under STIM1 deficiency induced by AβO and APOE4, and also evaluates the effects of STIM1 overexpression. Combined AβO and APOE4 treatment reduced cofilin Ser3 phosphorylation in neurons neurons relative to AβO co-administered APOE2 or APOE3 (Figure 4A). By contrast, phosphorylated LIM domain kinase (LIMK), the active form of the cofilin upstream kinase LIMK remained unchanged (Figure 4A). This finding implies that an alternative signaling pathway accounts for the observed differences in p-cofilin abundance.

**Figure 4.**
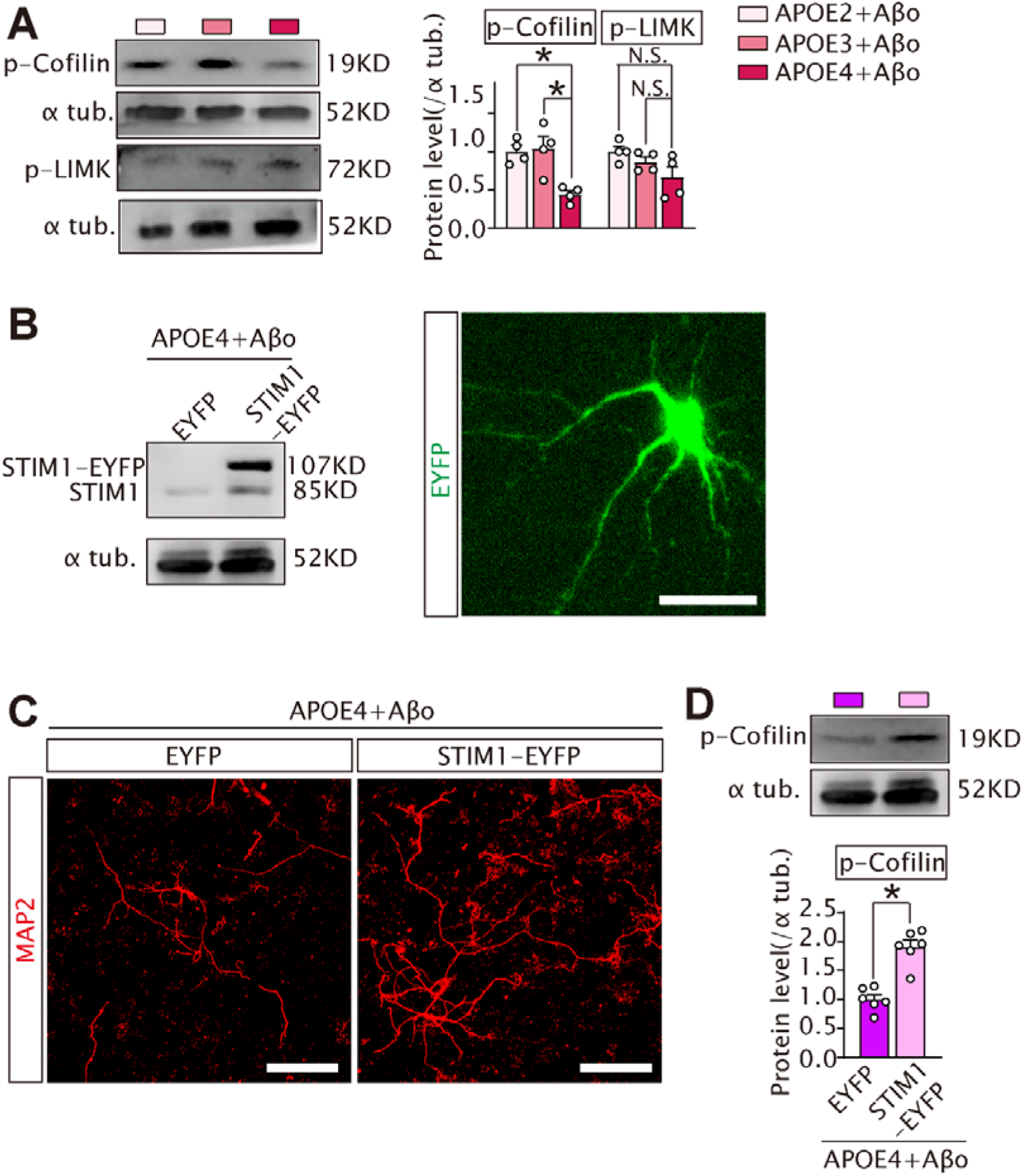
Overexpression of STIM1 restored dendritic arbors in hippocampal neurons under the treatments of APOE4 and AβO. Primary hippocampal neuronal cultures at DIV7-9 were subjected to treatments with AβO (200 nM), in conjunction with either APOE2, APOE3, or APOE4 (200 nM each) for 24 h. (**A**) Western blot was conducted to examine the levels of p-Cofilin (Ser3(1:1000)) and p-LIMK1 (Thr508)/LIMK2 (Thr505) (1:1000). One-way ANOVA followed by Bonferroni test was used for comparing the differences of p-Cofilin and p-LIMK. Four independent cell cultures were prepared for each treatment. p-Cofilin, F (2, 9) = 10.121, p =0.005, and p-LIMK, F (2, 9) = 3.089, p=0.095. The levels of p-Cofilin treated with APOE4 + AβO were either lower than those treated with APOE2 + AβO (p = 0.013), or lower than those treated with APOE3 + AβO (p = 0.009). The levels of p-LIMK in APOE4 + AβO treatment did not differ significantly from those either in APOE4 + AβO (p = 0.244), or in APOE3 + AβO group (p = 0.577). (**B**) Primary neuronal cultures at DIV4-5 were transfected with pEX-SP-EYFP-STIM1 (D76A) (STIM1-EYFP) or control plasmid, pEX-SP-EYFP (EYFP) for 48 h. STIM1 levels were shown in the transfected neurons (left panel). The expression was also observed under a Zeiss LSM880 confocal laser microscope system (Zeiss). Bar, 50μm. (**C**) After transfection, the neurons were treated with APOE4 plus AβO for 24 h. The neurons were fixed and stained for MAP2 (red). Images were obtained under 20×objective. Bar, 100μm for all panels. The images were representative of two slides of 12-13 neurons in either condition. (**D**) After transfection, the neurons were treated with APOE4 plus AβO for 24 h. p-Cofilin levels were assessed. A Student’s *t-test* was employed for comparison. The p-Cofilin in neurons transfected with STIM1-EYFP plasmid were significantly higher than those in neurons transfected with EYFP plasmid (n = 6 for each), indicating t(10) = 6.19, p <0.0001.

We further investigated whether the overexpression of STIM1 in neurons reversed dendritic arbor deficits treated with AβO and APOE4. Cultured hippocampal neurons were transfected with the pEX-SP-EYFP-STIM1 (D76A) plasmid to constitutively activate SOCE, following the method described in previous research ^22^. The levels of EYFP-STIM1 (D76A) protein in neurons transfected with the pEX-SP-EYFP-STIM1(D76A) plasmid were significantly higher than those in cells transfected with the control plasmid, pEX-SP-EYFP (Figure 4B). After transfection, the neurons were treated with AβO (200 nM) and APOE4 (200 nM) for 24 h. Compared to transfection with the control plasmid, dendritic arbors indicated by MAP2 staining exhibited longer dendrites and more branches when overexpressing EYFP-STIM1(D76A) (Figure 4C). The phosphorylation of cofilin at Ser3 in neurons overexpressing the active form of STIM1 was higher than neurons transfected with the control plasmid (Figure 4D).

### Combined AβO and APOE4 treatment activated Bcl2-dependent autophagic degradation of glutathione peroxidase 4

The results in Figure 2 strongly indicate that autophagy mediates the biological effects of combined AβO and APOE4 treatment. We next investigated whether combined treatment with AβO and APOE4 modulates autophagy in neurons. AβO combined with APOE4 decreased p62 compared to combined AβO and APOE3 or APOE2, while did not alter LC3BII levels (Figure 5A). Autophagy was further consolidated with transfection of a tandem mRFP-EGFP LC3 construct. The mRFP signal is more stable than the GFP signal in the acidic conditions of lysosomes. As expected, the treatment of AβO combined with APOE4 increased either red or merged puncta in neurons compared with the treatments of AβO combined with APOE2 (Figure 5B). Since lysosome is the final destination for autophagy influx, we examined the effect after pertubing lysosome function. Addition of NH_4_Cl, a chemical alkalizing lysosome, before the end of the treatment, increased the levels of LC3BII and p62 under combined AβO and APOE4 treatment (Figure 5C). Thus, co-treatment with AβO and APOE4 enhanced autophagy compared with co-administration of AβO and APOE2.

**Figure 5.**
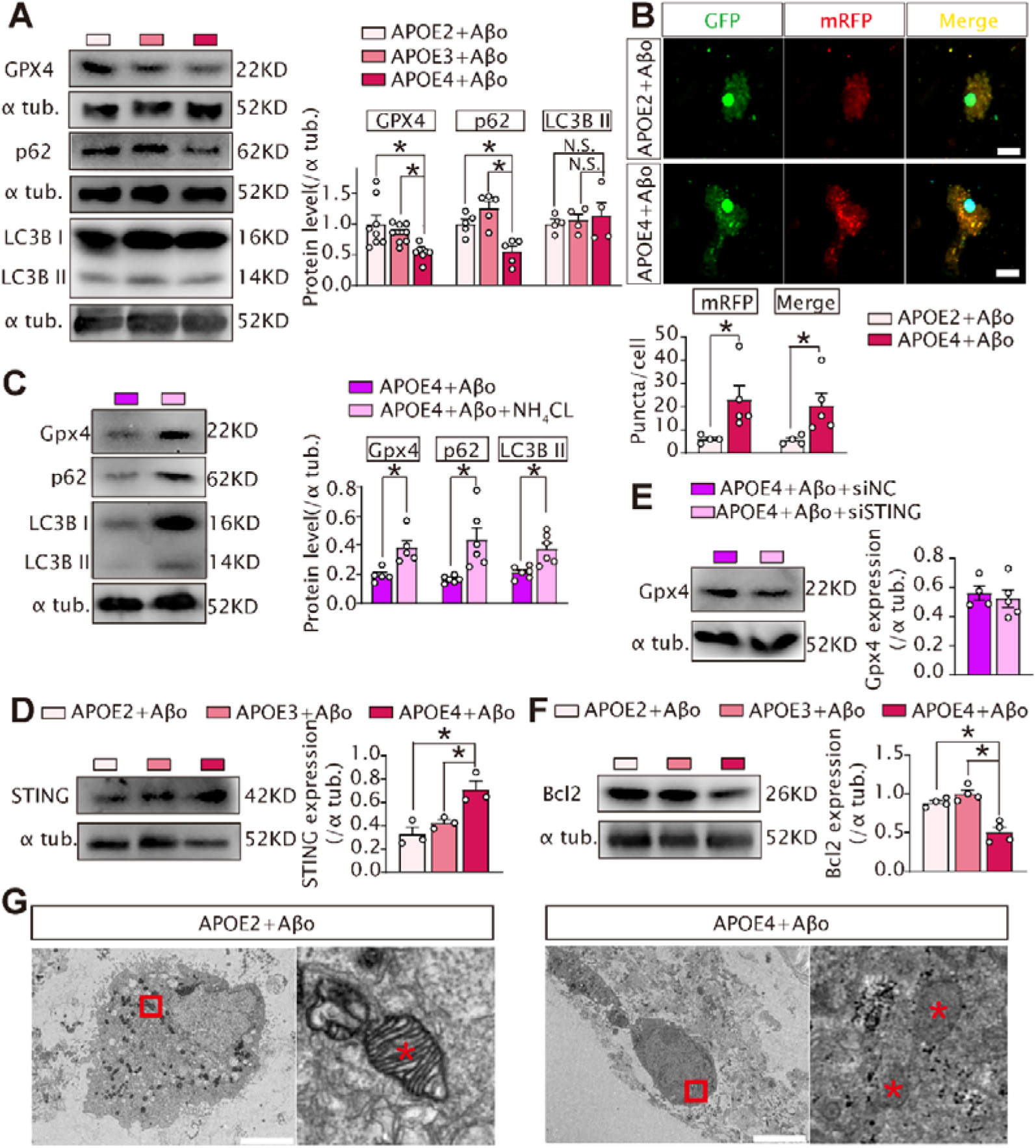
APOE4 together with AβO induced autophagy, which in turn degraded Gpx4. Primary neuronal cultures were subjected to treatments as indicated. (**A**) One-way ANOVA, followed by Tukey or Bonferroni test, was employed for comparing protein levels. Comparing Gpx4, F(2, 21) = 6.711, p =0.006. Gpx4 in APOE4 + AβO *versus* APOE2 + AβO (p = 0.039) or APOE3 + AβO (p = 0.002). Comparing LC3BII, F (2, 9) = 0.217, p = 0.809. p = 0.932, and 0.988 for thereof comparisons. Comparing p62, F (2, 12) = 13.634. p = 0.001. p = 0.019, and 0.001 for thereof comparison (n = 5-8 for each group). (**B**) Primary neuronal cultures were transfected with ptfLC3 plasmid and subsequently subjected to treatments as indicated. Cultures were observed under a confocal microscope, 63×objective. Bar, 10 μm. Red-only (mRFP) or merged puncta treated with APOE4+AβO were more than in APOE2 + AβO (n = 4-5) (Mann-Whitney U test, p= 0.016 for red-only and Student’s *t-test* for merged puncta comparison, t(7) = 2.44, p = 0.04). (**C**) The cultures were pre-treated with NH4Cl (10mM) or vehicle for 10 min, then treated with APOE4 plus AβO for 24 h. NH4Cl pre-treatment increased Gpx4, p62, or LC3BII levels. Student’s *t-test* was used for comparison (Gpx4, t(8) = 3.535, p = 0.008; p62, t(10) = 3.293, p = 0.021; LC3BII, t(10) = 3.524, p = 0.011), n = 5-6 for each group. (**D**) One-way ANOVA, followed by Tukey or Bonferroni test, was employed for comparing STING protein levels. STING levels in neurons treated with APOE4 + AβO were higher than those in APOE2 + AβO, or APOE3 + AβO-treated neurons, F(2, 6) = 11.755, p =0.008. APOE4 + AβO *versus* APOE2 + AβO, p = 0.010 or *versus* APOE3 + AβO (p = 0.039). (**E**) Primary neurons were transfected with STING-specific siRNA (siSting), or scrambled siRNA (siNC) with lipofectamine 3000^TM^. After 48 h following transfection, the neurons were treated with APOE4+ AβO for 24 h. Student’s *t-test* was used for comparing the differences of Gpx4 protein levels. Gpx4 levels in neurons transfected with the siSting (n = 4) were not statistically different from those transfected with siNC (n = 5), t(7) = 0.458, p=0.660. (**F**) One-way ANOVA, followed by Bonferroni test, was employed for comparing Bcl2 protein levels. Bcl2 proteins in neurons treated with APOE4 + AβO were lower than that in APOE2 + AβO, or APOE3 + AβO-treated neurons, F(2, 9) = 27.610, p< 0.0001. Bcl2 in APOE4 + AβO *versus* APOE2 + AβO (p = 0.001) or *versus* APOE3 + AβO (p< 0.0001). (**G**) Electron microscopic analysis was conducted to assess ultrastructure in neurons either treated with APOE2 + AβO or APOE4 + AβO. Representative images were obtained with transmission electron microscope system at a magnification of 2500. Red asterisks indicated mitochondria. Bar, 5μm.

Since glutathione peroxidase 4 (Gpx4) protects against ferropotosis by counteracting oxidative stress ^32^, we also assessed Gpx4 protein abundance. In neuronal cultures, co-treatment with AβO and APOE4 reduced Gpx4 relative to AβO co-administered with APOE3 or APOE2 (Figure 5A), consistent with elevated ROS triggered by AβO and APOE4. We next explored the mechanism underlying autophagy induction upon AβO and APOE4 co-treatment. A recent report showed that STIM1 downregulation drives STING-dependent autophagy in the context of multiple sclerosis ^33^. We thus tested whether STING mediates autophagy induction induced by AβO and APOE4 co-treatment. STING protein accumulated, whereas phosphorylated STING remained undetectable (Figure 5D). STING accumulation in the absence of p-STING indicates that STING was not activated, while ER-Golgi was disrupted. In line with this observation, knockdown of STING did not alter Gpx4 protein levels (Figure 5E). Attenuated SOCE has been shown to suppress calcium-calmodulin kinase activity ^18^, thereby blunting CREB activation. Moreover, CREB promotes transcription of the *Bcl2* gene ^34^. Bcl2 binds Beclin1 and blocks assembly of the autophagy initiation complex ^35^. We therefore examined Bcl2 protein under these treatment conditions. Co-treatment with AβO and APOE4 reduced Bcl2 abundance compared with combination of AβO and APOE3 or APOE2 (Figure 5F). Furthermore, electron microscopy revealed a marked reduction in mitochondrial cristae in neurons treated with AβO and APOE4 relative to neurons receiving AβO plus APOE2 (Figure 5G).

## Discussion

STIM1 localizes predominantly to the ER membrane and finely tunes calcium signaling between the ER calcium store and plasma-membrane calcium influx via ER calcium-dependent conformational changes ^36^. STIM1 activation is triggered upon calcium dissociation from its calcium-binding domains, which drives STIM1 oligomerization ^37^. This conformational rearrangement disrupts the intramolecular interaction between CRAC activation domain and the polybasic tail of STIM1, facilitating STIM1 binding to the ORAI channel and enabling calcium influx ^38, 39^. In addition, STIM1 exerts an inhibitory gating effect on the Ca_V_1.2 calcium channel ^20^. Reduced STIM1 expression may relieve this inhibition and thereby increase calcium influx, leading to calcineurin activation. Contrary to this observation, we detected no elevation in cytosolic calcium. We also did not observe nuclear translocation of TFEB (Supplemental Figure 2), a downstream readout governed by balance regulation of calcineurin and mTORC1. The VPS34 complex contributes to autophagy induction by facilitating autophagosome nucleation ^40^. Downregulation of Bcl2 upon combined AβO and APOE4 treatment disrupts Bcl2–Beclin1 interaction, freeing Beclin1 to assemble the VPS34 complex. Our findings therefore suggest that the VPS34 complex assembly, triggered by reduced Bcl2 abundance, mediates autophagy induction under cotreatment of APOE4 and AβO. Notably, another study reports that APOE4 suppresses autophagy by competing with TFEB for binding to the CLEAR motif within gene enhancer, thereby repressing transcription of autophagy-related genes ^41, 42^. This apparent discrepancy stems from differences in control treatments: our study employed APOE2 plus AβO as the control, whereas the referenced work used sham treatment.

Our previous study elucidated a mechanism whereby endoplasmic reticulum stress-induced STIM2 degradation simplifies dendritic arbors by altering the dynamics of phosphorylation and dephosphorylation of cofilin ^5^. In the present study, we also demonstrated that STIM1 depletion reduced p-cofilin, thereby influencing the actin dynamics, a major cytoskeletal component of dendritic spines. Notably, co-treatment with APOE4 and AβO led to the degradation of STIM1, but not STIM2, via p62-mediated selective autophagy. However, P62 expression was not upregulated but instead downregulated following 24 h of AβO and APOE4 exposure. We propose that oxidative stress induced by AβO and APOE4 triggers transient p62 accumulation at early timepoint, which is ultimately cleared through lysosomal targeting and degradation. Notably, STIM2 undergoes autophagic degradation under endoplasmic reticulum stress ^5^. APOE4 and AβO did not alter STIM2 abundance but downregulated STIM1, which may be attributed to their distinct subcellular distributions: STIM2 localizes to the ER membrane, while STIM1 is positioned adjacent to the plasma membrane.

Following STIM1 downregulation in neurons, we observed aberrent mitochondrial morphology marked by diminished mitochondrial cristae, implying that ferroptosis may accompany axonal degeneration. The mitochondrial morphological defects induced by STIM1 downregulation in our study align with a prior report demonstrating that STIM1 knockout in differentiated cells causes substantial impairment of mitochondrial function ^20^. A major limitation of this study is the absence of in vivo data from humanized APOE4 knock-in AD mouse models, as well as human post - mortem AD brain samples, which were not accessible to our laboratory. However, Gpx4 has been found to be downregulated in Alzheimer’s brain ^43^, which correlates with the accumulation of reactive oxygen species (ROS) characteristic of AD pathology. In addition, we have not established causal relationship between STIM1 reduction and decreased p-cofilin, given that p-LIMK levels remain unchanged under conditions of STIM1 downregulation. The Rac1-PAK-LIMK-cofilin signaling cascade promotes cofilin phosphorylation at Ser3, which inactivates cofilin and suppresses actin severing, ultimately stabilizing dendritic spines ^44^. Beyond this pathway, CaMKII and calcineurin also regulate cofilin phosphorylation ^45^. A prior study has shown that reduced Neuronal SOCE lowers CaMKII activity^18^. Therefore, cofilin dephosphorylation may be mediated by inhibited CaMKII under conditions of combined APOE4 and AβO treatment.

SOCE deficiency promotes neuronal loss by disrupting cytoplasmic and ER calcium homeostasis ^5, 18, 20, 46, 47^. Conversely, excessive SOCE activation is associated with neuronal loss driven by tau accumulation. Targeting septins can restore SOCE activity and thereby suppress tau aggregation ^48, 49^. Given the distinct temporal windows of Aβ and tau pathologies ^1^, SOCE may exert stage-specific effects in AD via divergent mechanisms. This heterogeneity in AD progression should be considered when developing therapeutic strategies to modulate SOCE.

C1q-triggered classical complement pathway activation mediates synapse elimination during physiological development and in the AD brain, with microglia serving as key mediators in this process ^6, 50^. Of note, the present study uncovers an novel intrinsic neuronal mechanism driving synapse elimination that operates independently of microglial involvement. Notably, all experiments in this study were performed using purified neuronal cultures, which excludes microglial interference.

In summary, we demonstrate that AβO and the APOE4 isoform impair SOCE by downregulating STIM1. STIM1 overexpression rescues deficits in dendritic arborization induced by AβO and the APOE4. Mechanistically, autophagy-mediated degradation of Gpx4, subsequent ROS accumulation, and disrupted mitochondrial morphology constitute downstream consequences of STIM1 reduction driven by AβO and the APOE4. Collectively, we conclude that AβO and the APOE4 suppress STIM1 expression, which in turn drives autophagic degradation of the antioxidant protein Gpx4, further promoting ROS buildup and mitochondrial dysfunction. Our findings uncover a novel mechanism underlying APOE4-potentiated AβO synaptotoxicity.

## Declarations

### Ethics approval and consent to participate

All experimental procedures following Animal Research Reporting of the *In Vivo* Experiments (ARRIVE) Guidelines and the protocols used were also approved by the Wenzhou Medical University Committee on the Use and Care of Animals (wydw2020-0056). All mice were provided with water and food *ad libitum* in a standard pathogen-free facility with automatic illumination on a 12-hour on/off cycle at Wenzhou Medical University.

### Data availability statement

All data are contained in the main text and in supplemented figures. The original data including quantifications and the statistics in excel form have been deposited in public data repository in China, SciencDB, at the doi link, 10.57760/sciencedb.38372.

## Acknowledgements

We appreciate financial support of the above funding agency.

## Author contributions

Zhou J, investigation, images acquisition, and writing; Liao HJ, investigation; Liang WJ, investigation; Liu SY, investigation; Cai J, materials; Liu Y, investigation; Liu YM, investigation; Zhou ZL, materials; Wu S, conceptualization, supervision, and writing

## Abbreviations

AD: Alzheimer’s disease
APOE: apolipoprotein E
SOCE: Store-operated calcium entry
STIM: stromal interaction molecule
Gpx4: glutathione peroxidase 4
MAP2: microtubule-associated protein 2
Nrf2: Nuclear factor erythroid 2-related factor 2
ARE: antioxidant response element
ROS: reactive oxygen species
TFEB: transcription factor EB

