## Supplementary files for "Apolipoprotein E4 Exacerbates Aβ Oligomer-Induced Synaptotoxicity through Promoting Autophagic Degradation of Glutathione Peroxidase 4"

**Supplemental Figures**

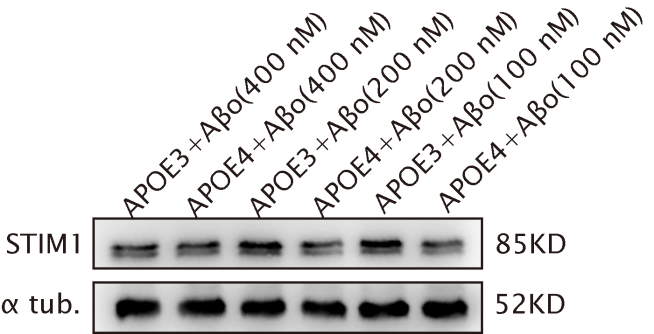

**Supplemental Figure 1. Effects of co-administeration of different concentrations**

**of AβO and APOE4 on STIM1.** Primary hippocampal neuronal cultures at days *in vitro* (DIV) 7-9 were treated with AβO (100, 200, 400 nM), and along with APOE3, or 4 (100, 200, 400 nM each) for 24 h. Western blot was conducted to analyze STIM1 levels in the protein lysates.

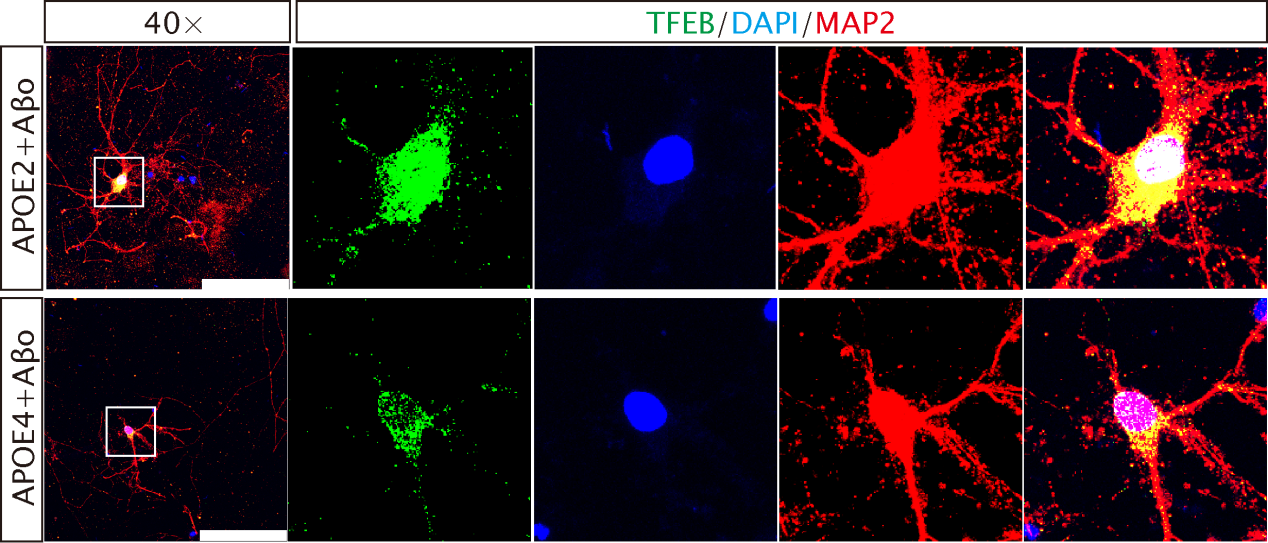

**Supplemental Figure 2. AβO and APOE4 did not induce TFEB nuclear translocation, relative to AβO and APOE2.** MAP2 (red) and TFEB (green), and DAPI (blue) were co-stained in neurons treated with APOE2 (200 nM) + Aβo (200 nM) or APOE4 (200 nM) + Aβo (200 nM) for 24 h. APOE4+Aβo (14 neurons, 3 dishes) and APOE2+Aβo (12 neurons, 3 dishes). Images were obtained with Leica stellaris5 system under 40× objective, Bar, 100μm for all panels.
